# Optogenetic stimulation of memory-tagged neurons elicits endogenous patterns of neural activity

**DOI:** 10.1101/2023.05.15.540888

**Authors:** Claudia Jou, José R. Hurtado, Danielle Cruz-Holden, Simón Carrillo-Segura, Eun Hye Park, André A. Fenton

## Abstract

Hippocampal neurons encoding memories discharge in sparsely-distributed patterns; they can be optogenetically tagged, then photostimulated to elicit conditioned behavior, potentially generating a synthetic memory trace, “engram”. We investigated mouse hippocampal population responses to the photostimulation of memory-tagged neurons to determine if memory-associated discharge is mimicked in hippocampus, a network with non-linear and homeostatic interactions. Both memory-tagged and not-tagged CA1 cells adjusted firing during photostimulation without altering place cell firing fields. Cell-pair cofiring relationships also maintain during photostimulation, indicating a low-dimensional, dynamical structure-preserving, homeostatic network response instead of the photostimulated pattern. Photostimulating neurons that were tagged during place-avoidance memory elicits similar place-avoidance memory in control conditions. Thus artificial photostimulation elicits natural, stored, homeostatic neuronal network cofiring patterns to elicit memory, establishing mimicry evidence for the engram.

## Introduction

The mouse hippocampus is a neuronal system that is crucial for memory and other cognitive functions like navigation. Unlike sensory neurons that respond to stimuli, the firing rates and cofiring statistics of hippocampal neurons encode abstract cognitive variables like location, and are substantially internally generated, and interdependent with external stimuli ^1-3^. Importantly, minute-scale hippocampal population activity is homeostatic across environments and diverse behaviors, including sleep ^4,5^. In contrast, individual neuronal components of the hippocampal network change characteristically to encode specific items of information, like events ^6,7^ and locations in the case of hippocampal place cells ^8^. This collection of adaptive, non-linear responses and interactions result from feedforward and feedback inhibition, characteristic of an inhibition stabilized network. Accordingly, the hippocampus is an archetype of a complex system, making the network effects of cellular manipulations difficult to interpret ^9-12^.

### A neural network model illustrating that periodic optogenetic stimulation of a neuron subset can produce homeostatic instead of periodic network responses

Optogenetics, chemogenetics, and other neuron-specific manipulations have been revolutionary by enabling so-called causal experiments that selectively manipulate specific neurons to evaluate their hypothesized function. Despite enabling an unprecedented ability to identify functional neuronal circuits by the effective connections amongst neurons, it remains unclear how to correctly interpret the results of these precise manipulations of a system with adaptive, non-linear, and homeostatic network interactions (Fig. 1A). Specifically, we seek to understand how optogenetic stimulation of memory-associated neurons can elicit the expression of the conditioned response as has been demonstrated in so-called “engram cell” studies ^13,14^. We examine the effects of photostimulation of neurons that were optogenetically tagged during behavioral tasks, by evaluating the responses of individual cells as well as the collective neuronal network responses ^15,16^. While it is possible to manipulate and measure a change in a specific component of a neuronal circuit (Fig. 1A), the consequence of that same manipulation can have hard to predict network-wide effects that cannot be evaluated by measuring isolated component changes (Fig. 1B), in part because they dynamically depend on network correlations and other features ^17,18^. It is even possible that a manipulation elicits changes throughout the individual system components that are collectively homeostatic, resulting in the system not changing, especially in sparse inhomogeneous networks, which have very few opportunities for controllability ^19^, in accord with Ashby’s law of requisite variety ^20^. Accordingly, “homeostatic” network hypotheses predict that in an engram cell experiment, photostimulation will elicit familiar network-wide intrinsically homeostatic responses, whereas “causal” hypotheses predict novel stimulation-controlled network responses.

**Figure 1.**
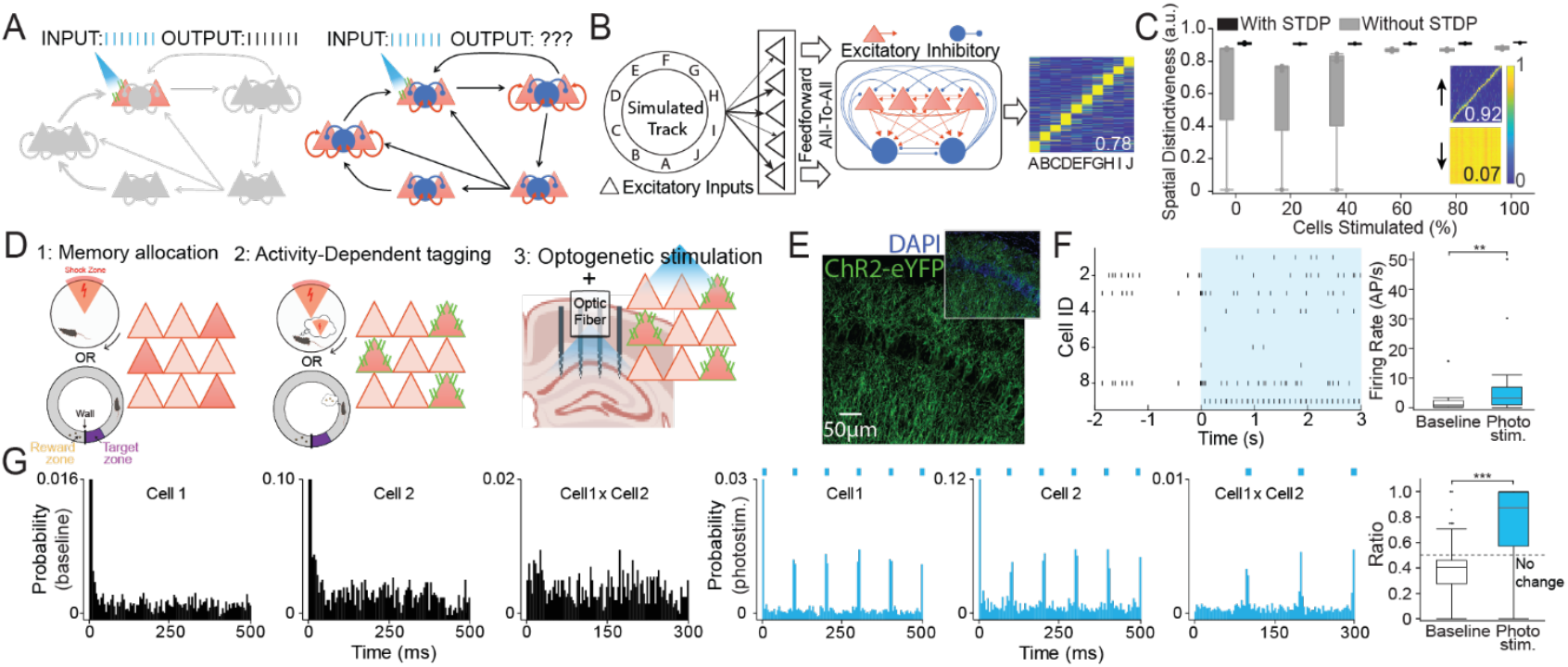
Optogenetic tagging and photostimulation of neural circuits *in vivo*. Optogenetically-tagged neurons might respond predictably to photostimulation as if A) network effects were minimal or B) respond unpredictably due to strong network effects. C) Neural network architecture illustrating that the 10 locations (A-J) of a ring-shaped environment are randomly mapped to “sensory” units which in turn project to a network of 500 excitatory (E) and 50 inhibitory (I) units that are interconnected by activity dependent STDP-modifiable E-E, E-I, and I-I connections. With experience the E units acquire location-specific activity patterns, here shown as a matrix with x-axis: locations, y-axis: cell identity sorted by location of peak activity, and average activity indicated by the blue-to-yellow color scale. The location-specificity is quantified as spatial distinctiveness. C) Spatial distinctiveness is minimally changed by periodically stimulating a random proportion of E cells, demonstrating that optogenetic stimulation could have minimal effects on network activity. D) Schematic experimental design illustrates memory training and optogenetic tagging during either i) an active place avoidance task followed by ensemble recordings and photostimulation either during anesthesia or head-fixation or ii) during shuttling on a circular track for food reward followed by ensemble recordings and photostimulation during the shuttling task. E) Immunofluorescence identified memory-tagged CA1 neurons. F) Under urethane anesthesia, electrophysiological effects of 10-Hz photostimulation of place avoidance memory-tagged CA1 principal cells. A raster and summary firing rate statistics from ensemble recordings before and during photostimulation are shown, as well as G) auto- and cross-correlograms of two neurons during (left) baseline and (middle) photostimulation; (right) summary autocorrelation change ratios. ^*^p<0.05, ^**^p<0.01, ^***^p<0.001.

To gain intuition for how it is possible to activate components of a network without changing the network’s behavior, we began by using a heuristic neural network model of excitatory and inhibitory cells in which the excitatory cells learn to distinguish the locations along a circular track with location-specific firing because of spike timing-dependent plasticity (STDP) rules Fig. 1C; ^21,22^. After sufficient training, excitatory photostimulation was simulated by periodically activating a random subset of the excitatory cells during exploration of the track. Consistent with homeostasis, but potentially contradicting causal interpretations of key experiments ^23,24^, stimulation of up to 30% of the model cells only rarely (20%) disturbed location-specificity. Even stimulating 40% or more sometimes did not disturb location-specificity (Fig. 1D); the results were similar with the STDP rules off during stimulation (Fig. 1D, S1), motivating us to investigate whether or not (causal) stimulus-induced or (homeostatic) intrinsic network patterns are elicited by optogenetic stimulation of neurons in the intact mouse brain.

### Photostimulation of memory-tagged CA1 neurons causes reliable action potential responses under anesthesia

During the expression of a long-term active place avoidance memory, the mostly excitatory neurons that (strongly) expressed the immediate early gene *Arc* were indelibly tagged with an enhanced yellow fluorescent protein (eYFP) and channelrhodopsin (ChR2) fusion protein, consequent to transient Cre-mediated recombination Figs. 1E,F and S2; ^25,26,27^. Weeks later, under urethane anesthesia, the ChR2-tagged hippocampal CA1 neurons were activated by 10-Hz light pulses that increased firing (Fig. 1G), as predicted by causal hypotheses. Both auto- and cross-correlograms show 10-Hz 473-nm photostimulated neuronal activation (Fig. 1G; t_25,52_ = 4.43, p = 0.0002). Remarkably, 45% of recorded cells increased 10-Hz periodicity in their autocorrelation despite only 15% were immunohistochemically verified to be tagged (test of proportions z = 8.84, p < 0.0001; Fig. S2) suggesting synchronous network activation by photostimulation under anesthesia.

### Photostimulation of memory-tagged CA1 neurons causes homeostatic network responses in awake behaving mice

The response to photostimulation appeared more complex in drug-free awake behaving mice (Fig. 2A). The light pulses evoked responses in some but not all cells (Fig. 2B); 18.6% of cells were considered optotagged in that they expressed an evoked response (Fig. 2C), similar to the (15%) immunohistological confirmation of tagging (test of proportions z = 1.63, p = 0.1; Fig. S2). Mice received either 4-Hz (n = 3) or 10-Hz (n = 6) stimulation, while they were head-fixed (n = 6 mice, 210 cells) or shuttling for food reward on a circular track (n = 4 mice, 515 cells, Fig. S4), with similar effects unless indicated otherwise. We considered two network models that predict distinct steady-state responses to photostimulation (Fig. 2E). The “engram cell” model assumes photostimulation is reliably both effective and selective for tagged cells, leaving not-tagged cells undisturbed (Fig. 2E, F top). The “homeostatic” model assumes network effects of photostimulation dominate, causing similarly modest activation of tagged and non-tagged cells and elevated interneuron firing due to network-level E-I coupling and homeostasis (Fig. 2E, F bottom). The observed responses to photostimulation during head-fixation and shuttling were steady after the first minute (Fig. 2G). During head-fixation, 67% of tagged cells increased firing compared to 32% of not-tagged cells (z = 3.42, p < 0.0001). During shuttling behavior, 22% of tagged and 19% of not-tagged cells increased firing (z = 0.60, p = 0.54). Forty-four presumptive inhibitory interneurons were recorded in the two conditions; two of these increased firing, indistinguishable from the expectation of zero since they are not tagged (test of proportions z = 1.34, p = 0.18). Despite constant photostimulation, in both head-fixed and the shuttling conditions, these increases appeared greater during the first 30 s of stimulation and decreased to steady-state values by 2 min, irrespective of whether or not the cells were tagged but this observation was not statistically robust (30 s vs. 120 s: z’s ≤ 1.73, p’s ≥ 0.08).

**Figure 2.**
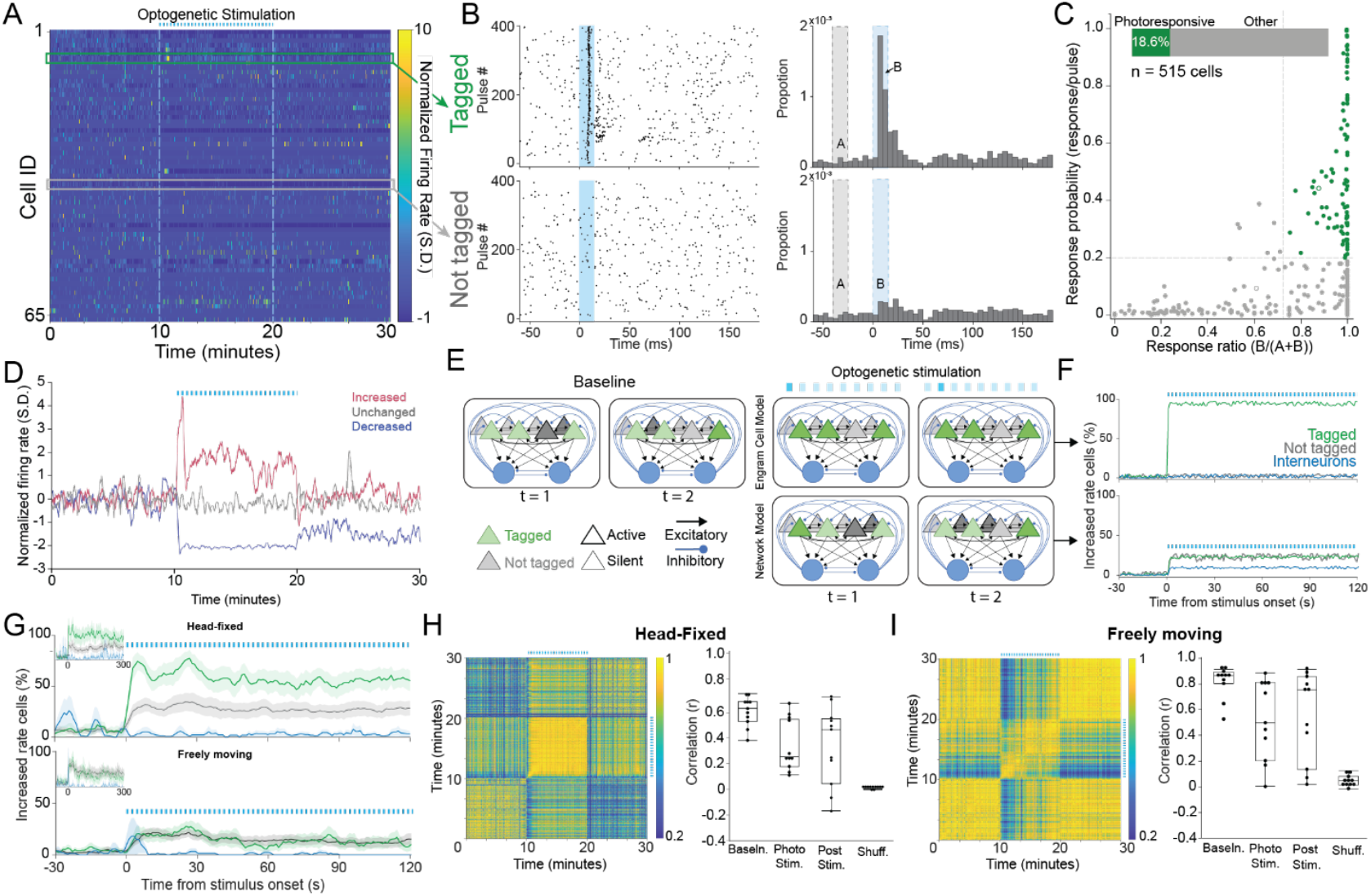
Optogenetic stimulation elicits complex, endogenous patterns of CA1 discharge in the awake behaving mouse. A) Color-coded activity of a 65-cell CA1 ensemble recorded from a freely-behaving mouse before, during and after 4-Hz photostimulation. Optogenetic tagging and the recordings were during the shuttling task. B) Peristimulus time raster (left) and histogram (right) of the initial 400 stimulus pulses of a tagged and a not-tagged cell and C) the quantitatively classified photoresponsivity of 515 CA1 cells (of the 675 cells identified during baseline 515 were followed during photostimulation). The 96 (18.6%) that were photoresponsive are classified as tagged (green). D) Firing rate time series of three example cells before, during and after 4-Hz photostimulation depict diverse and dynamic responses to photostimulation. (E) Pictorial depiction of the engram cell and network models and F) their predictions of photoresponsivity, compared to G) observed photoresponsivity. H) Head-fixed and I) freely-behaving mouse examples of a (left) correlation matrix characterizing similarity of 10-s ensemble activity vectors before, during and after photostimulation and (right) summary quantification of the similarity to baseline activity. Data from 6 head-fixed and 4 freely-behaving mice.

The overall pattern of network discharge assessed as 10-s ensemble activity vector correlations was reliable (greater than shuffled) compared to baseline, but more stable during baseline than during photostimulation in both head-fixed (Fig. 2H; F_2,30_ = 5.93, p < 0.0001, post hoc tests baseline > photostim = poststim) and shuttling behaviors, consistent with the observed rate changes (Fig. 2I; F_2,27_ = 3.37, p = 0.05, post hoc tests baseline > photostim = poststim). These observations were similar for 1-s activity vectors (Fig. S3) and confirm that optogenetic stimulation elicits a homeostatic network response that resembles before as well as poststimulation, despite individual neurons firing faster and slower. Instead of persistently controlling network discharge, photostimulation generates a complex homeostatic network response. Subsequent analyses excluded the first minute of photostimulation to better assess the steady-state effect of stimulation.

### CA1 place cell responses are stable during photostimulation of CA1 memory-tagged neurons in freely behaving mice

We then investigated place cell discharge during the shuttling behavior to compare predictions of the engram cell and homeostatic hypotheses (Fig. 3A). Engram cell hypotheses predict that place cell tuning is disrupted by periodic photostimulation that controls neuronal discharge of cells that were tagged during the same behavior in the same environment (Fig. 3B top). In contrast, homeostatic hypotheses predict place cell tuning maintains during the photostimulation (Fig. 3B bottom). Mice learn to shuttle in the circular track for a food reward, a behavior that is not affected by photostimulation (Fig. 3C and Fig. S5). Relative to pre-stimulation baseline, place fields appear normal and unchanged during and after photostimulation (Fig. 3D). Place and non-place cells had similar probabilities of being tagged (Fig. 3E; n = 79 place cells, n = 255 non-place cells; test of proportions z = 0.85, p = 0.39), the proportions of place cells observed during and after photostimulation are also not different from baseline (Fig. 3F; test of proportions z’s ≤ 1.40, p’s ≥ 0.16). The quality of place fields also does not differ between the tagged and non-tagged cells as measured by place information content (Fig. 3G) and coherence (Fig. 3H; Information content: F_1,154_ = 0.59, p = 0.4; Coherence: F_1,154_ = 0.11, p = 0.7), nor before, during, or after photostimulation (Information content: F_2,154_ = 1.03, p = 0.4; interaction: F_2,154_ = 0.04, p = 0.9; Coherence: F_2,154_ = 2.05, p = 0.1; interaction: F_2,154_ = 0.59, p = 0.6). We conclude that during photostimulation the stability of place fields is indistinguishable from during baseline (Fig. 3I).

**Figure 3.**
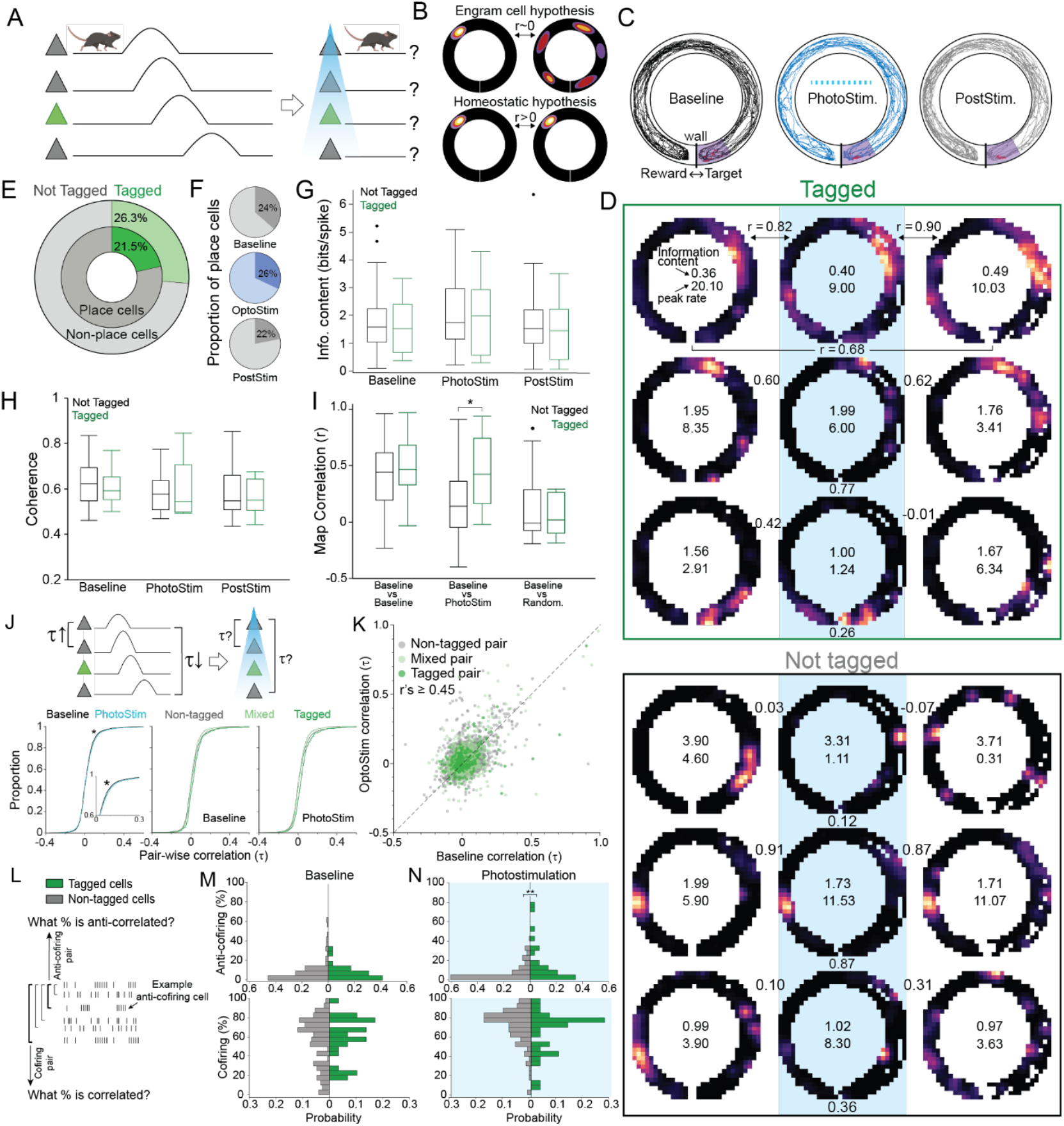
Photostimulation of tagged cells does not change CA1 place cell firing fields. A) Two predictions for how causal optogenetic photostimulation might affect place cell firing fields during shuttling on a linear track. B) Engram cell hypotheses predict disrupted firing fields in tagged neurons that respond to periodic stimulation that is indifferent to current position, whereas homeostatic hypotheses predict stable firing fields. C) Example spatial tracks, and D) firing rate maps during recordings of tagged and not-tagged place cells during baseline, photostimulation and post-stimulation. E) Overall and F) session-specific likelihoods of spatial firing classifications. Session-specific spatial tuning quality compared between tagged and not-tagged cells measured by G) spatial information content, and H) coherence, as well as their I) spatial firing stability relative to baseline. J) Measuring how photostimulation affects cofiring during shuttling. Baseline and photostimulation distributions of correlation (t) measures of cofiring comparing tagged cell pairs (dark green, n = 279 pairs), mixed tagged and not-tagged cell pairs (light green, n = 1809 pairs) and non-tagged cell pairs (gray, n = 3372 pairs). K) Scatterplot comparing the cofiring of cell pairs before (Baseline) and during photostimulation. L) The anti-cofiring power and the cofiring power was determined for each cell. Histograms show the likelihood that cells participate in (upper) anti-cofiring and (lower) cofiring pairs (M) before and (N) during photostimulation in the head-fixed mice. ^*^p<0.05, ^**^p<0.01, ^***^p<0.001.

### Stable cofiring and evoked environment-specific anti-cofiring during photostimulation of CA1 memory-tagged neurons in freely behaving mice

We next examine the cell-pair cofiring (τ) statistics at 1-s time resolution (Fig. 3J) to estimate the seconds-scale network response ^21,28^, which should be disrupted according to the engram cell hypothesis and maintained according to the homeostatic hypothesis. Correlations measuring cofiring are stronger during compared to before photostimulation (Fig. 3K; F_1,11971_ = 7.75, p = 0.005). Consistent with tagged and not-tagged cells responding distinctly to photostimulation, cofiring during photostimulation was greater if either both cells in a pair or none in the pair were tagged, compared to if only one cell of the pair was tagged (F_2,11971_ = 36.00, p < 0.0001; interaction F_1,11971_ = 0.36, p = 0.69; Tukey tests both = none > either). Nonetheless, the cofiring of individual cell pairs was stable between baseline and photostimulation (Fig. 3M; df = 5182, r = 0.48, p < 0.0001) whether both (df = 274, r = 0.54, p < 0.0001), one (df = 1744 r = 0.52, p < 0.0001), or none of the cells in a pair was tagged (df = 3160, r = 0.47, p < 0.0001). These relationships were similar when the timescale to assess cofiring was varied from 40 ms to 5 s (Fig. S6).

Of the baseline cofiring relationships, 9% were significantly negative (Fig. 3L, “anti-cofiring pairs”) and 70% were significantly positive (“cofiring pairs”). Because individual CA1 cells that have many anti-cofiring relationships with other cells are environment-specific and crucial for discriminating environmental representations ^21^, we examined whether the tagged and non-tagged cells differed in their anti-cofiring power. The mice that were tagged during place avoidance were recorded during head-fixation, so we examined whether photostimulation causes them to anti-cofire within the population. While the tagged and non-tagged cells in these mice did not differ at baseline (Fig. 3M; t_181_ = 0.63, p = 0.3), the tagged cells were more likely to be anti-cofiring with other cells during photostimulation (Fig. 3N; t_181_ = 2.28, p = 0.002). Correspondingly, tagged cells were less likely than non-tagged cells to cofire with other cells during photostimulation (t_181_ = 1.86, p = 0.03). These differences in cofiring relationships were not evident in the shuttling mice that were tagged and recorded during shuttling in the same environment (Fig. S5). Indeed, consistent with the homeostatic hypothesis, these distinct anti-cofiring responses to photostimulation are predicted by the network model which also suggests the homeostatic responses depend on synaptic plasticity (Fig. S6).

### Ensemble activity manifold structure and neurocentric coordinates during photostimulation of CA1 memory-tagged neurons in freely behaving mice

Projecting the 1-s high-dimensional neural activity vectors into a 3-D subspace using the non-linear IsoMap dimensionality reduction algorithm preserves high-dimensional neighborliness, allowing us to define and visualize low-dimensional structure (“topology”) and coordinates (“geometry”) of the high-order cofiring relationships within neuronal ensemble activity Fig. 4A; ^29^. We were motivated to examine the ensemble population dynamics because prior work shows that 15-20% of the most anti-cofiring subpopulation of CA1 cells is responsible for distinguishing manifold representations of distinct environments ^21^. Accordingly, photostimulating tagged neurons, which preserves most cofiring and may selectively increase anti-cofiring of tagged cells (Fig. 3N), might register the homeostatic topology of ensemble activity to the current environment, potentially explaining how stimulation of memory-tagged cells can elicit the conditioned behavior that was expressed during tagging ^3,13,14,21,22^. The baseline and photostimulated manifold projections were similar (Fig. 4B-D). The centroids of photostimulated activity only differed from baseline in the primary dimension (Fig. 4E; F_1,174_ = 13.76, p = 10^-4^) indicating photostimulation shifts the coordinates of the population geometry consistent with the registration expectation (Fig. 4F)^3^. Other measures of population geometry and topology did not differ between baseline and photostimulation (Fig. 4G,H), and the overlap of the two low-dimensional projections was indistinguishable from the overlap of two halves of the baseline recording (Fig. 4I). Properties of the time-dependent trajectories (distance and localness) through the 3-D subspace were similar during baseline and photostimulation, whereas smoothness was reduced during photostimulation (t_17_ = 6.47, p = 10^-6^; Fig. 4J-M), consistent with a homeostatic response that maintains cofiring (Fig. 3M), despite the photostimulation potentially activating the environment-specific anti-cofiring subset that shifts the population activity’s geometry and potential registration to the environment ^3,21,30,31^. These findings indicate that the optogenetic stimulation was not sufficient to qualitatively change the neurocentric organization of CA1 ensemble activity, and that photostimulation may register the internally-organized representation of the tagged experience to the current environment, consistent with homeostatic hypotheses and evidence ^3,32^.

**Figure 4.**
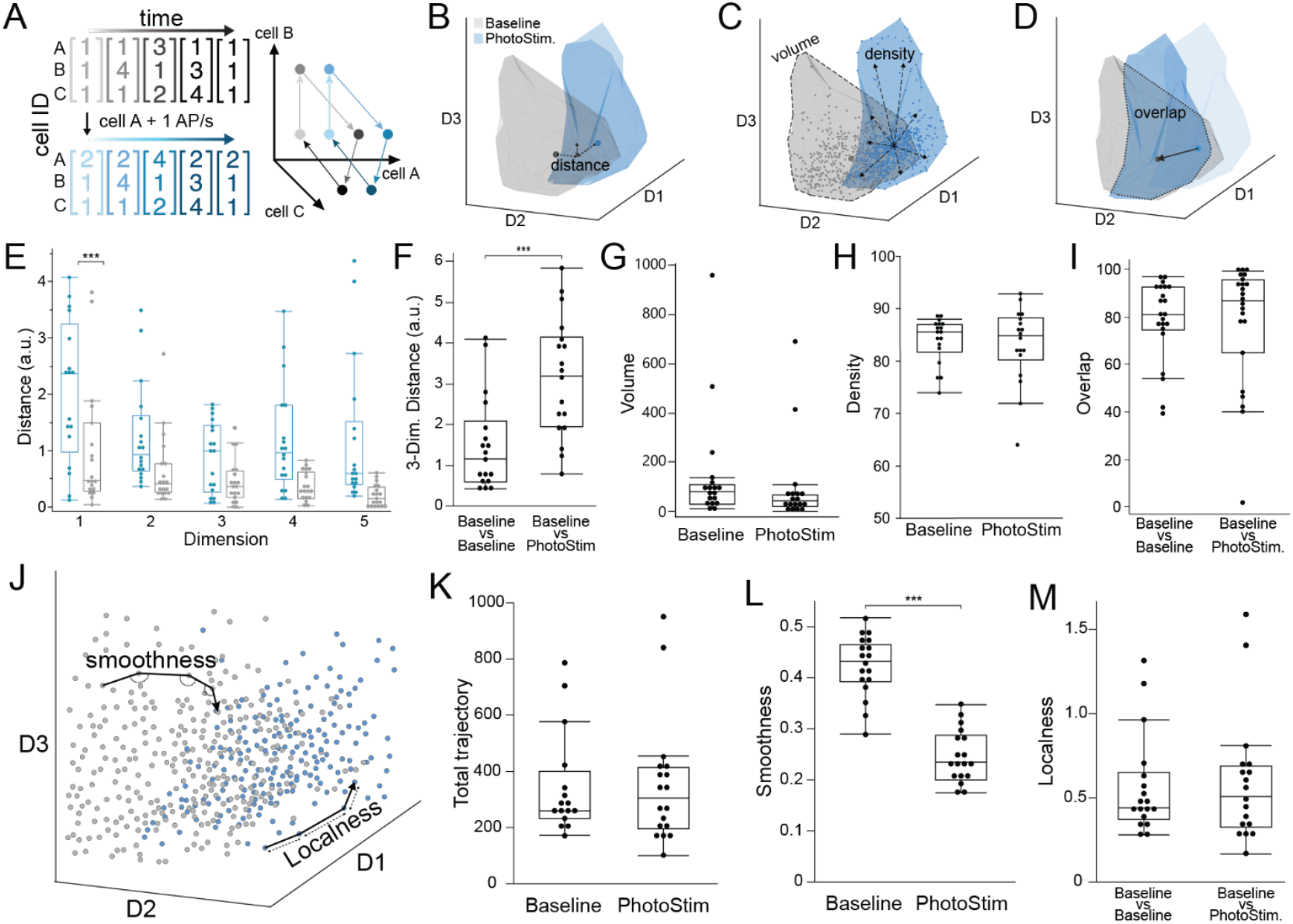
Optogenetic stimulation shifts the low-dimensional population geometry of CA1 ensemble discharge. A) The ensemble activity of n cells is observed in a n-dimensional state space, which non-independent cofiring organizes on compact manifolds in low-dimensional subspaces. The manifold structure (“topology”) is preserved when cofiring relationships are maintained and the neurocentric coordinates in state space (“geometry”) shifts when activity changes along one or more subspace dimensions. B-D) The 3-D ensemble activity of an example 26-cell recording before and during photostimulation projected into a 3-D IsoMap subspace. Each 3-D projection illustrates different measures of the 3-D embedding. E) Distance between baseline and photostimulated manifold centroids in each of the top-five IsoMap dimensions and F) the Euclidean distance in the top-three dimensions. Comparisons of baseline and photostimulated manifold measures of G) volume, and H) density, and I) overlap with the baseline manifold. J) The trajectory of ensemble activity through the 3-D state space compared during baseline and photostimulation by measures of the trajectory K) distance, L) smoothness, and M) localness. ^*^p<0.05, ^**^p<0.01, ^***^p<0.001.

### Optogenetic photostimulation of CA1 memory-tagged neurons elicits a conditioned active avoidance

Is the CA1 network’s relative indifference to the photostimulation just the property of a complex system, is the photostimulation sufficient to elicit the memory that was tagged, or is the photostimulation simply insufficient to be consequential? To decide between these alternatives, we examined whether the optogenetic photostimulation was sufficient to elicit the conditioned active place avoidance behavior in an environment where the mice do not express an avoidance. Mice received 10 minutes of active place avoidance training three times a day for four days in one environment. Each day, before or after one of the place avoidance trials, the mice were put for 10 minutes in a separate neutral environment that was identical except that transparent plastic covered the floor, the ambient scent was different, and shock was never on (Fig. 5A). After four days of training the mice did not express an avoidance in the neutral environment and Arc-CreER^T2^-eYFP-ChR2 experimental mice and Arc-CreER^T2^-eYFP control mice received 4OH-Tamoxifen (TAM) to tag active neurons. Seven Arc-CreER^T2^-eYFP-ChR2 (“memory-ChR2”) mice and four Arc-CreER^T2^-eYFP (“memory-eYFP”) mice received TAM before being put in the place avoidance environment. Six Arc-CreER^T2^-eYFP-ChR2 (“neutral-ChR2”) mice received TAM before being put in the neutral environment. Two weeks later, all mice were returned to the neutral environment and as in all previous experiments, photostimulated at 473-nm and either 4-Hz (n = 8) or 10-Hz (n = 9) with similar effects. Whereas memory-eYFP and neutral-ChR2 mice walked throughout the neutral environment, memory-ChR2 mice tended to avoid an area corresponding to a shock zone (Fig. 5B), which could be described for each mouse by the Rayleigh vector of the dwell angles within the circular environment (Fig. 5C), as well as thigmotaxis that is characteristic of training sessions. Avoidance was quantified by the probability of being in the corresponding shock zone, analyzed in two 5-min epochs. The probability was decreased in the memory-ChR2 mice (Fig. 5D; F_2,27.8_ = 10.3, p = 0.0005), and increased in the second epoch (F_1,27.8_ = 5.3, p = 0.03), with no group X epoch interaction (F_2,27.8_ = 0.44, p = 0.6; Tukey tests: memory-ChR2 < neutral-ChR2 = memory-eYFP). Similarly, the memory-ChR2 mice avoided the zone for longer periods of time than the neutral-ChR2 and memory-eYFP mice (Fig. 5D; F_2,26.6_ = 6.8, p = 0.004; Tukey tests: memory-ChR2 > neutral-ChR2 = memory-eYFP). Neither the effects of epoch nor the group X epoch interaction were significant (F’s ≤ 0.9, p’s ≥ 0.3). These group differences were not due to the mice walking at different speeds (F_2,28_ = 1.7, p = 0.2).

**Figure 5.**
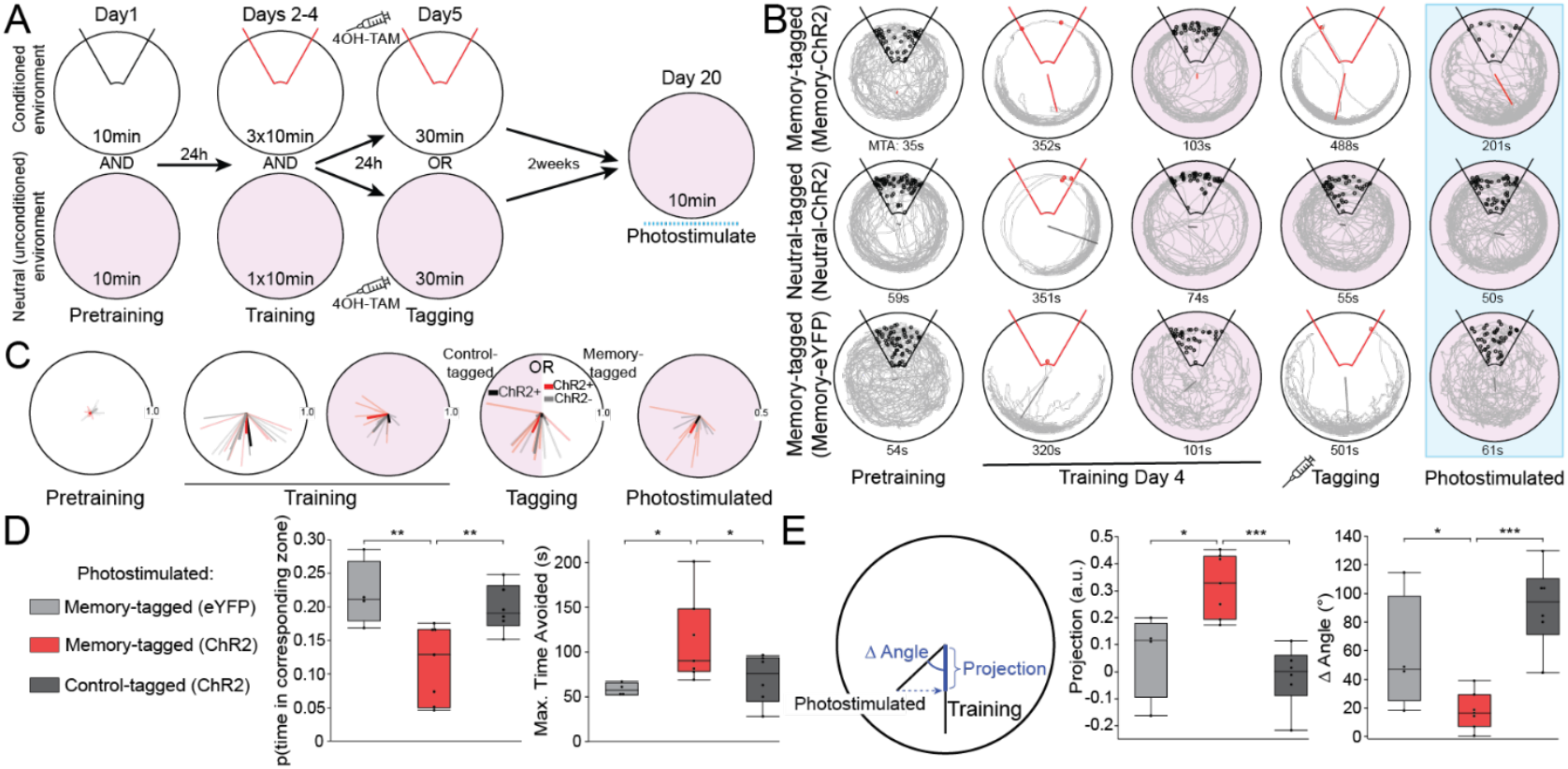
Optogenetic stimulation elicits conditioned active place avoidance behavior in the unconditioned environment, despite never experiencing shock there. A) Experimental design. B) Tracks of three example mice in the conditioned and unconditioned environments during select behavioral trials. Each resultant vector overlay indicates the preferred polar location. C) Distribution of resultant vectors across the behavioral protocol. Color-code as in panel D. Avoiding the location in the unconditioned environment that corresponds to the shock zone in the conditioned environment quantified as D) the probability of being in the zone (left) and MTA, the maximum time of avoiding the zone (right). E) The similarity of the photostimulated avoidance to the shock-elicited avoidance was quantified by the projection of the light-elicited resultant vector onto the training-elicited resultant vector (left) and summarized for each group by the distributions of the projection length (middle) and angle (right). ^*^p<0.05, ^**^p<0.01, ^***^p<0.001.

Because each mouse’s place avoidance was idiosyncratic in the conditioned environment, avoidance during the last training session on day 5 was characterized by the Rayleigh vector of all the mouse’s angular positions. These tended to be long and idiosyncratically oriented away from the shock zone for avoidance training sessions and short for corresponding sessions on the neutral environment (Fig. 5C). If photostimulation in the neutral environment causes the mice to express their particular avoidance, then each mouse’s Rayleigh vector during photostimulation in the neutral environment should resemble the mouse’s Rayleigh vector during the last training session. The length of the projection of the “photostimulation” vector onto the “training” vector was longest in the memory-ChR2 group (Fig. 5E; F_2,14_ = 12.74, p = 0.001, Tukey tests: memory-ChR2 > neutral-ChR2 = memory-eYFP). The angular difference between the photostimulation and training vectors were also smallest in the memory-ChR2 group (Fig. 5E; F_2,14_ = 11.96, p = 0.001, Tukey tests: memory-ChR2 < memory-eYFP = neutral-ChR2). These findings provide compelling evidence that the photostimulation of memory-ChR2 mice in the neutral environment is sufficient to elicit the specific learned avoidance that had been tagged during the last training session before photostimulation.

## Discussion

Photostimulating a subset of active place avoidance memory-tagged cells is sufficient to elicit behavioral expression of the memory in a familiar neutral environment, yet the same stimulation did not elicit novel patterns of neural discharge in the hippocampus output (see Video S1 for photostimulation that elicits the conditioned avoidance in a memory-ChR2 mouse but not a neutral-ChR2 mouse, in a completely novel environment). Memory-tagging as performed here was driven by *Arc* activation that only captures a subset of memory-activated cells that likely only weakly overlap with cells that activate immediate early genes *Fos* and *Npas4* that are each sufficient to identify hippocampal cells that are crucial for an aversive memory ^27,33^ . Although their activity is sufficient for eliciting the conditioned response, it is unclear whether the memory-tagged cells are the most Arc-activated and/or the most electrophysiologically active subpopulation ^34,35^. Our experimental design extends the “engram cell” reactivation phenomenon from non-spatial threat avoidance conditioning ^13,14^, demonstrating that optogenetic tagging and photostimulation is also effective for an active place avoidance memory that is exquisitely sensitive to hippocampal function ^36,37^ and dependent on the months-long persistence of PKMζ-maintained long-term potentiation in hippocampus ^26,38,39^. As is the case for threat avoidance memories, we expect that active place avoidance memories activate neurons across distributed brain regions, not just hippocampus ^40^. Engram cell activation experiments tend to photostimulate for 3-6 minutes. We observed that after an initial, transient activation trend lasting < 30 s, the CA1 network adapts to continued photostimulation (Figs. 2,3), consistent with homeostatic hypotheses and incompatible with causal hypotheses. Prudence is thus warranted when interpreting the results of experimental designs that intend causal manipulations of neuronal circuits in awake, behaving animals, because as demonstrated, the manipulations elicit homeostatic, endogenous patterns of activity rather than the stimulus-triggered responses that are more likely under anesthesia, a condition that may compromise excitation-inhibition coordination Fig. 1; ^41^. While feedforward conceptions of neural circuits predict linear responses to tagged-neuron photostimulation as demonstrated in reduced preparations ^42^, we demonstrate substantially invariant network responses despite single neuron activity changes, as expected if feedback, recurrence, inhibition, and other neural connectivity motifs provide for complex system dynamics Fig. 1; ^19,43^. While these motifs include and depend upon synaptic function, some engram cell experiments have been interpreted to indicate that modified synaptic function has no role in memory storage, which is at odds with the present findings ^24,44^. We conclude that unless neural activity is measured directly, the outcomes of engram-cell, behavioral timescale synaptic plasticity, and similar neural system manipulations should be parsimoniously interpreted as the expression of the homeostatic, endogenous population dynamics of a complex system, rather than the precise causal responses of a neuronal circuit.

## Acknowledgements

Many thanks to György Buzsáki for critical comments on earlier drafts of the manuscript. We are especially grateful for the encouragement from Gyuri and Mauricio Rangel-Gomez. Supported by NIH grant R01MH132204.

## Funding

Supported by National Institute of Health grant R01MH132204.

## Author Contributions

Conceptualization: CJ and AF

Methodology: CJ, DH-C and EHP

Investigation: CJ, JH, DH-C and EHP

Software: CJ, JH, SS-C, and AF

Visualization: CJ and JH

Funding acquisition: AF

Project administration: CJ and AF

Supervision: AF

Writing – original draft: CJ, JH, and AF

Writing – review & editing: CJ, JH, DC-H, AF

## Inclusion and ethics

This research was conducted with respect for the animal subjects, the individual researchers and their unique perspectives, and our collective effort to learn and generate individually useful knowledge.

## Competing interests

The authors declare no competing interests.

## Data availability

The datasets are available from the corresponding author upon reasonable request. Source data are available at <u>data repository</u>

## Code availability

Code used for data processing and analysis is available from the corresponding author upon reasonable request at <u>code repository</u>

## Notes

### Competing Interest Statement

The authors have declared no competing interest.

### Summary of Updates

This revision includes analyses of place cell firing during shuttling behavior within a circular track. CA1 place cell action potential discharge of individual cells as well as the patterns of firing across the population are compared before and during photostimulation. Additional modeling is also performed to demonstrate feasibility of the observations within he theoretical context of a neuronal network population doctrine rather than the framework of the standard neuron doctrine.

## References

1. Stark, E., Roux, L., Eichler, R., and Buzsaki, G. (2015). Local generation of multineuronal spike sequences in the hippocampal CA1 region. Proc Natl Acad Sci U S A 112, 10521–10526. 10.1073/pnas.1508785112.

2. Dragoi, G., and Tonegawa, S. (2013). Distinct preplay of multiple novel spatial experiences in the rat. Proc Natl Acad Sci U S A 110, 9100–9105. 1306031110 [pii] 10.1073/pnas.1306031110.

3. Fenton, A.A. (2024). Remapping revisited: how the hippocampus represents different spaces. Nature Reviews Neuroscience. 10.1038/s41583-024-00817-x.

4. Buzsaki, G., Csicsvari, J., Dragoi, G., Harris, K., Henze, D., and Hirase, H. (2002). Homeostatic maintenance of neuronal excitability by burst discharges in vivo. Cereb Cortex 12, 893–899.

5. Dragoi, G., Harris, K.D., and Buzsaki, G. (2003). Place representation within hippocampal networks is modified by long-term potentiation. Neuron 39, 843–853. S0896627303004653 [pii].

6. Lenck-Santini, P.P., Fenton, A.A., and Muller, R.U. (2008). Discharge properties of hippocampal neurons during performance of a jump avoidance task. J Neurosci 28, 6773–6786. 28/27/6773 [pii] 10.1523/JNEUROSCI.5329-07.2008.

7. Moita, M.A., Rosis, S., Zhou, Y., LeDoux, J.E., and Blair, H.T. (2003). Hippocampal place cells acquire location-specific responses to the conditioned stimulus during auditory fear conditioning. Neuron 37, 485–497. S0896627303000333 [pii].

8. O’Keefe, J. (1976). Place units in the hippocampus of the freely moving rat. Exp Neurol 51, 78–109. 0014-4886(76)90055-8 [pii].

9. Watkins de Jong, L., Nejad, M.M., Yoon, E., Cheng, S., and Diba, K. (2023). Optogenetics reveals paradoxical network stabilizations in hippocampal CA1 and CA3. Current Biology 33, 1689-1703.e1685. 10.1016/j.cub.2023.03.032.

10. Sanzeni, A., Akitake, B., Goldbach, H.C., Leedy, C.E., Brunel, N., and Histed, M.H. (2020). Inhibition stabilization is a widespread property of cortical networks. Elife 9. 10.7554/eLife.54875.

11. Rogers, S., Rozman, P.A., Valero, M., Doyle, W.K., and Buzsaki, G. (2021). Mechanisms and plasticity of chemogenically induced interneuronal suppression of principal cells. Proc Natl Acad Sci U S A 118. 10.1073/pnas.2014157118.

12. Dubreuil, A., Valente, A., Beiran, M., Mastrogiuseppe, F., and Ostojic, S. (2022). The role of population structure in computations through neural dynamics. Nature Neuroscience 25, 783–794. 10.1038/s41593-022-01088-4.

13. Ramirez, S., Liu, X., Lin, P.A., Suh, J., Pignatelli, M., Redondo, R.L., Ryan, T.J., and Tonegawa, S. (2013). Creating a false memory in the hippocampus. Science 341, 387–391. 341/6144/387 [pii] 10.1126/science.1239073.

14. Garner, A.R., Rowland, D.C., Hwang, S.Y., Baumgaertel, K., Roth, B.L., Kentros, C., and Mayford, M. (2012). Generation of a synthetic memory trace. Science 335, 1513–1516. 335/6075/1513 [pii] 10.1126/science.1214985.

15. Vyas, S., Golub, M.D., Sussillo, D., and Shenoy, K.V. (2020). Computation Through Neural Population Dynamics. Annual Review of Neuroscience 43, 249–275. 10.1146/annurev-neuro-092619-094115.

16. Park, E.H., Keeley, S., Savin, C., Ranck, J.B., Jr., and Fenton, A.A. (2019). How the Internally Organized Direction Sense Is Used to Navigate. Neuron 101, 1–9. 10.1016/j.neuron.2018.11.019.

17. Posfai, M., Liu, Y.Y., Slotine, J.J., and Barabasi, A.L. (2013). Effect of correlations on network controllability. Sci Rep 3, 1067. 10.1038/srep01067.

18. Barabasi, D.L., Bianconi, G., Bullmore, E., Burgess, M., Chung, S., Eliassi-Rad, T., George, D., Kovacs, I.A., Makse, H., Papadimitriou, C., et al. (2023). Neuroscience needs Network Science. ArXiv.

19. Liu, Y.-Y., Slotine, J.-J., and Barabási, A.-L. (2011). Controllability of complex networks. Nature 473, 167–173. 10.1038/nature10011.

20. Ashby, W.R. (1958). Requisite variety and its implications for the control of complex systems. Cybernetica 1, 83–99.

21. Levy, E.R.J., Carrillo-Segura, S., Park, E.H., Redman, W.T., Hurtado, J.R., Chung, S., and Fenton, A.A. (2023). A manifold neural population code for space in hippocampal coactivity dynamics independent of place fields. Cell Reports 42, 113142. 10.1016/j.celrep.2023.113142.

22. Hurtado, J.R., Chung, S., and Fenton, A.A. (2025). Effective computations for hippocampal place cell phenomena in sparse untrained random networks. bioRxiv, 2025.2010.2004.680487. 10.1101/2025.10.04.680487.

23. Tonegawa, S., Liu, X., Ramirez, S., and Redondo, R. (2015). Memory Engram Cells Have Come of Age. Neuron 87, 918–931. 10.1016/j.neuron.2015.08.002.

24. Tonegawa, S., Pignatelli, M., Roy, D.S., and Ryan, T.J. (2015). Memory engram storage and retrieval. Curr Opin Neurobiol 35, 101–109. 10.1016/j.conb.2015.07.009.

25. Vazdarjanova, A., Ramirez-Amaya, V., Insel, N., Plummer, T.K., Rosi, S., Chowdhury, S., Mikhael, D., Worley, P.F., Guzowski, J.F., and Barnes, C.A. (2006). Spatial exploration induces ARC, a plasticity-related immediate-early gene, only in calcium/calmodulin-dependent protein kinase II-positive principal excitatory and inhibitory neurons of the rat forebrain. J Comp Neurol 498, 317–329. 10.1002/cne.21003.

26. Hsieh, C., Tsokas, P., Grau-Perales, A., Lesburgueres, E., Bukai, J., Khanna, K., Chorny, J., Chung, A., Jou, C., Burghardt, N.S., et al. (2021). Persistent increases of PKMzeta in memory-activated neurons trace LTP maintenance during spatial long-term memory storage. Eur J Neurosci 54, 6795–6814. 10.1111/ejn.15137.

27. Denny, C.A., Kheirbek, M.A., Alba, E.L., Tanaka, K.F., Brachman, R.A., Laughman, K.B., Tomm, N.K., Turi, G.F., Losonczy, A., and Hen, R. (2014). Hippocampal memory traces are differentially modulated by experience, time, and adult neurogenesis. Neuron 83, 189–201. S0896-6273(14)00404-8 [pii] 10.1016/j.neuron.2014.05.018.

28. Schneidman, E., Berry, M.J., 2nd, Segev, R., and Bialek, W. (2006). Weak pairwise correlations imply strongly correlated network states in a neural population. Nature 440, 1007–1012. nature04701 [pii] 10.1038/nature04701.

29. Tenenbaum, J.B., de Silva, V., and Langford, J.C. (2000). A global geometric framework for nonlinear dimensionality reduction. Science 290, 2319–2323. 10.1126/science.290.5500.2319.

30. Dragoi, G. (2024). The generative grammar of the brain: a critique of internally generated representations. Nat Rev Neurosci 25, 60–75. 10.1038/s41583-023-00763-0.

31. Buzsaki, G., McKenzie, S., and Davachi, L. (2022). Neurophysiology of Remembering. Annu Rev Psychol 73, 187–215. 10.1146/annurev-psych-021721-110002.

32. McKenzie, S., Huszar, R., English, D.F., Kim, K., Christensen, F., Yoon, E., and Buzsaki, G. (2021). Preexisting hippocampal network dynamics constrain optogenetically induced place fields. Neuron 109, 1040–1054 e1047. 10.1016/j.neuron.2021.01.011.

33. Sun, X., Bernstein, M.J., Meng, M., Rao, S., Sorensen, A.T., Yao, L., Zhang, X., Anikeeva, P.O., and Lin, Y. (2020). Functionally Distinct Neuronal Ensembles within the Memory Engram. Cell 181, 410–423 e417. 10.1016/j.cell.2020.02.055.

34. Guzowski, J.F., Miyashita, T., Chawla, M.K., Sanderson, J., Maes, L.I., Houston, F.P., Lipa, P., McNaughton, B.L., Worley, P.F., and Barnes, C.A. (2006). Recent behavioral history modifies coupling between cell activity and Arc gene transcription in hippocampal CA1 neurons. Proc Natl Acad Sci U S A 103, 1077–1082. 0505519103 [pii] 10.1073/pnas.0505519103.

35. Mizuseki, K., and Buzsaki, G. (2013). Preconfigured, skewed distribution of firing rates in the hippocampus and entorhinal cortex. Cell Rep 4, 1010–1021. S2211-1247(13)00401-4 [pii] 10.1016/j.celrep.2013.07.039.

36. Cimadevilla, J.M., Fenton, A.A., and Bures, J. (2000). Functional inactivation of dorsal hippocampus impairs active place avoidance in rats. Neurosci Lett 285, 53–56. S0304-3940(00)01019-3 [pii].

37. Cimadevilla, J.M., Wesierska, M., Fenton, A.A., and Bures, J. (2001). Inactivating one hippocampus impairs avoidance of a stable room-defined place during dissociation of arena cues from room cues by rotation of the arena. Proc Natl Acad Sci U S A 98, 3531–3536. 10.1073/pnas.051628398 98/6/3531 [pii].

38. Pastalkova, E., Serrano, P., Pinkhasova, D., Wallace, E., Fenton, A.A., and Sacktor, T.C. (2006). Storage of spatial information by the maintenance mechanism of LTP. Science 313, 1141–1144. 313/5790/1141 [pii] 10.1126/science.1128657.

39. Pavlowsky, A., Wallace, E., Fenton, A.A., and Alarcon, J.M. (2017). Persistent modifications of hippocampal synaptic function during remote spatial memory. Neurobiol Learn Mem 138, 182–197. 10.1016/j.nlm.2016.08.015.

40. Roy, D.S., Park, Y.G., Kim, M.E., Zhang, Y., Ogawa, S.K., DiNapoli, N., Gu, X., Cho, J.H., Choi, H., Kamentsky, L., et al. (2022). Brain-wide mapping reveals that engrams for a single memory are distributed across multiple brain regions. Nat Commun 13, 1799. 10.1038/s41467-022-29384-4.

41. Chung, A., Jou, C., Grau-Perales, A., Levy, E.R.J., Dvorak, D., Hussain, N., and Fenton, A.A. (2021). Cognitive control persistently enhances hippocampal information processing. Nature 600, 484–488. 10.1038/s41586-021-04070-5.

42. Boyden, E.S., Zhang, F., Bamberg, E., Nagel, G., and Deisseroth, K. (2005). Millisecond-timescale, genetically targeted optical control of neural activity. Nat Neurosci 8, 1263–1268. 10.1038/nn1525.

43. Liu, Y.-Y., and Barabási, A.-L. (2016). Control principles of complex systems. Reviews of Modern Physics 88, 035006. 10.1103/RevModPhys.88.035006.

44. Ryan, T.J., Roy, D.S., Pignatelli, M., Arons, A., and Tonegawa, S. (2015). Memory. Engram cells retain memory under retrograde amnesia. Science 348, 1007–1013. 10.1126/science.aaa5542.

45. Srinivas, S., Watanabe, T., Lin, C.S., William, C.M., Tanabe, Y., Jessell, T.M., and Costantini, F. (2001). Cre reporter strains produced by targeted insertion of EYFP and ECFP into the ROSA26 locus. BMC Dev Biol 1, 4. 10.1186/1471-213x-1-4.

46. Lacagnina, A.F., Brockway, E.T., Crovetti, C.R., Shue, F., McCarty, M.J., Sattler, K.P., Lim, S.C., Santos, S.L., Denny, C.A., and Drew, M.R. (2019). Distinct hippocampal engrams control extinction and relapse of fear memory. Nat Neurosci 22, 753–761. 10.1038/s41593-019-0361-z.

47. Perusini, J.N., Cajigas, S.A., Cohensedgh, O., Lim, S.C., Pavlova, I.P., Donaldson, Z.R., and Denny, C.A. (2017). Optogenetic stimulation of dentate gyrus engrams restores memory in Alzheimer’s disease mice. Hippocampus 27, 1110–1122. 10.1002/hipo.22756.

48. Neymotin, S.A., Lytton, W.W., Olypher, A.V., and Fenton, A.A. (2011). Measuring the Quality of Neuronal Identification in Ensemble Recordings. J Neurosci 31, 16398–16409. 31/45/16398 [pii] 10.1523/JNEUROSCI.4053-11.2011.

49. Pachitariu, M., Steinmetz, N., Kadir, S., Carandini, M., and Harris, K.D. (2016). Kilosort: realtime spike-sorting for extracellular electrophysiology with hundreds of channels. bioRxiv 061481 (2016). z10.1101/061481.

50. Levy, E.R.J., Carrillo-Segura, S., Park, E.H., Redman, W.T., Hurtado, J., Chung, S., and Fenton, A.A. (2021). A manifold neural population code for space in hippocampal coactivity dynamics independent of place fields. bioRxiv, 2021.2007.2026.453856. 10.1101/2021.07.26.453856.

