## Supplementary Information and figures for "Optogenetic stimulation of memory-tagged neurons elicits endogenous patterns of neural activity"

SUPPLEMENTARY MATERIAL

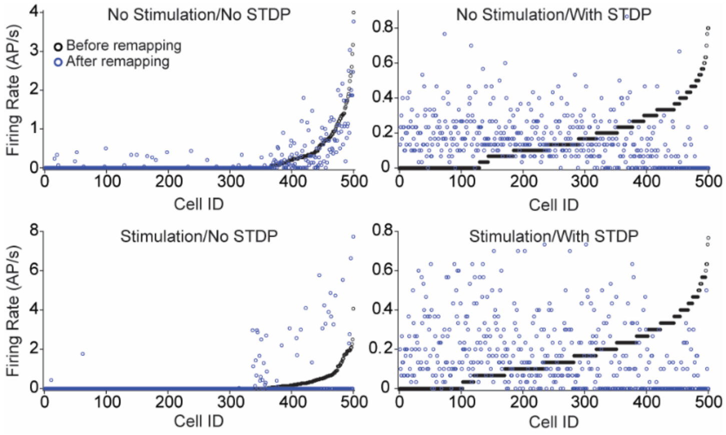


Figure S1. (related to Figure 1). Heuristic model effects of how stimulating excitatory units can change firing rates when sensory inputs from a novel ring environment are remapped into the winner-take-all competitive neural network of 500 excitatory (E) and 50 inhibitory (I) units in Figure 1, that has already learned the locations of a different ring environment under spike-timing dependent plasticity (STDP) learning rules. This simulation was performed to gain an intuition for the possible effects of optogenetic stimulation of engram cells in a different environment. The plots present average E unit activity rates before and after remapping, sorted from low-to-high-rate units. A random 20% of E units were stimulated in the lower plots to simulate optogenetic stimulation. While the effects of the stimulation are modest on rates, remapping strongly changes E rates only if STDP remains turned on; the rates are only weakly increased by stimulation if STDP is turned off.

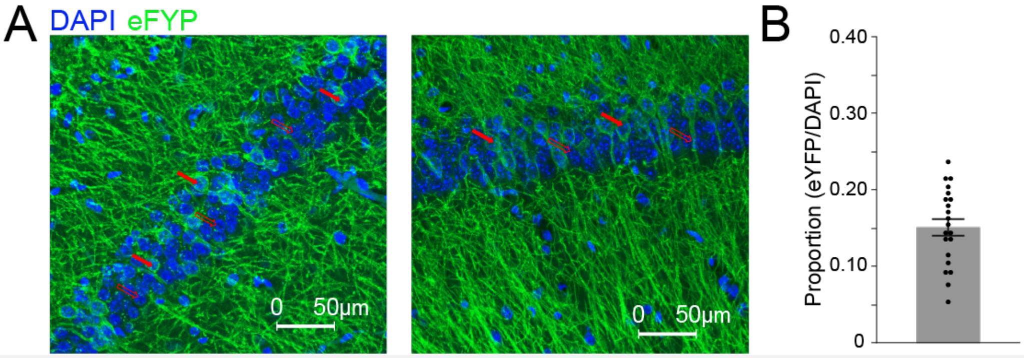


Figure S2 (related to Figure 1). Immunohistochemistry for DAPI (blue) and eYFP (green) to quantify the proportion of memory-tagged cells in CA1. A) Two example sections. B) DAPI-labeled CA1 pyramidal cell bodies were manually counted and the subset that were eYFP-positive established the proportion (15.1 ± 5%) of memory-tagged cells in each section (n = 3 sections per mouse, n = 7). Filled and open arrows show examples of tagged and not-tagged cells, respectively. Mean ± SEM.

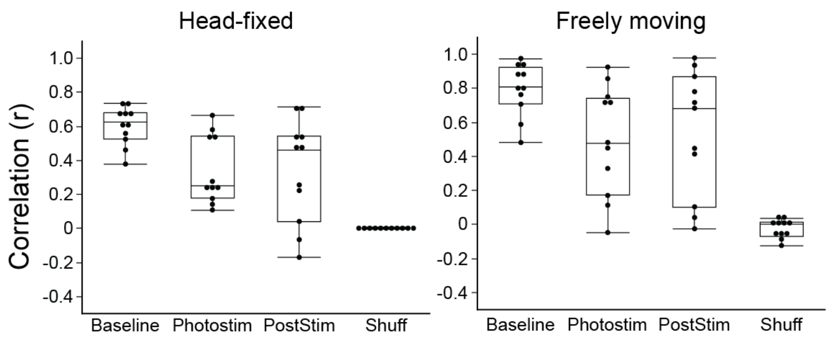


Figure S3 (related to Figure 2). Maintained activity vectors during photostimulation. Similarity of 1-s ensemble activity vectors before, during and after photostimulation compared to baseline from head-fixed (n = 6) and freely-behaving mice (n = 4). Each point is a single recording.

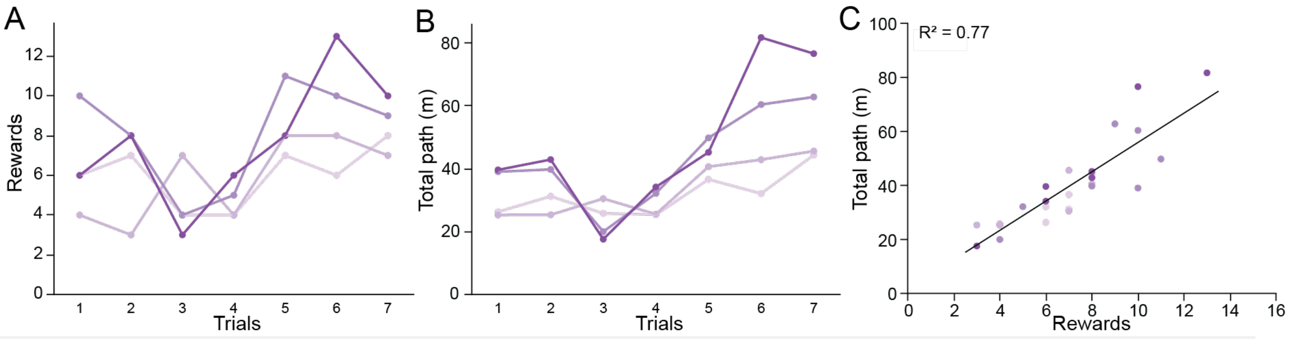


Figure S4. Measures of shuttling performance across daily trials. A) Rewards obtained, B) Total path covered, and C) their relationship.

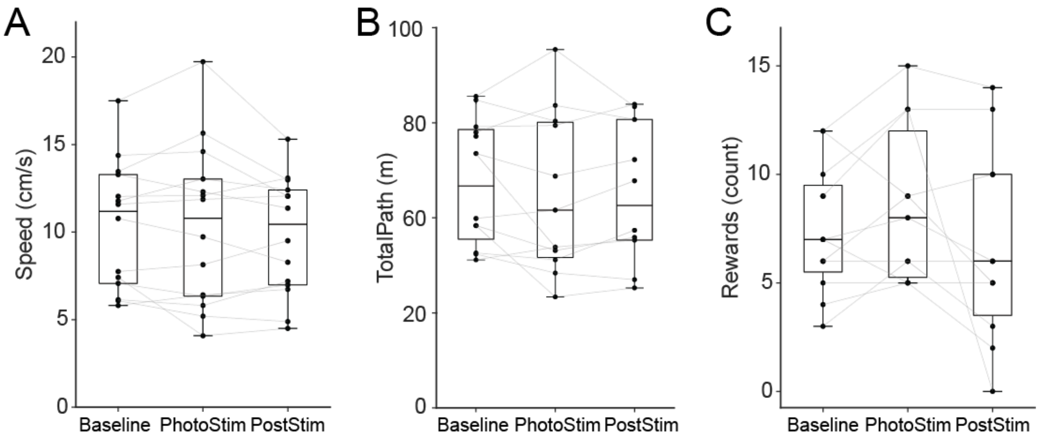


Figure S5. Summary statistics of shuttling behavior in 14 sessions of 4 mice before, during and after photostimulation of neurons that were tagged during the shuttling task

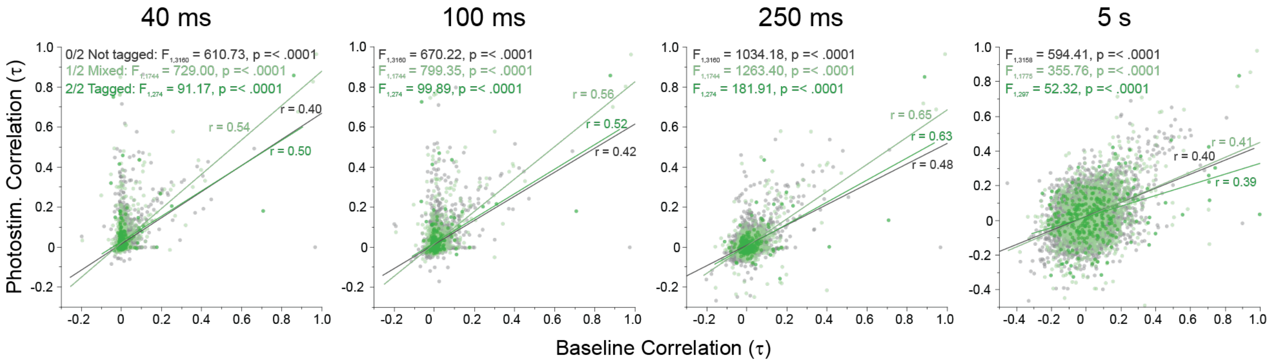


Figure S5 (related to Figure 3). Comparing cofiring during baseline and photostimulation in the different pairs of tagged and not-tagged cells at the different timescales indicated above the plots.

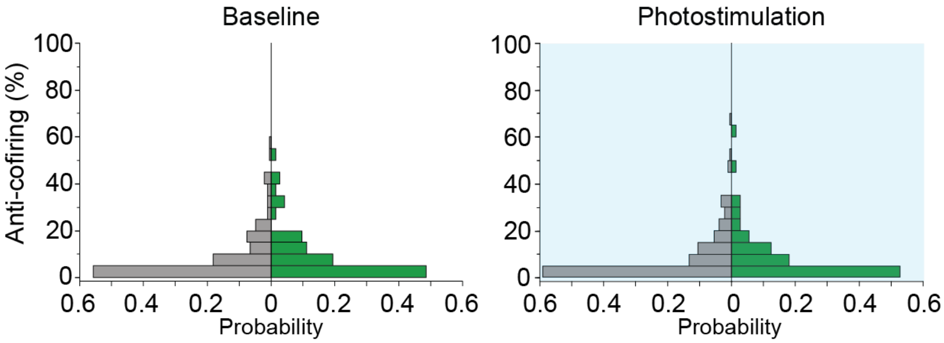


Figure S6 (related to Figure 3). Anti-cofiring in shuttling mice is unchanged by photostimulation. The percentage of cell pairs in which a particular cell was significantly negatively correlated ("anti-cofiring") was determined for each cell. Histograms show the likelihood that cells participate in anti-cofiring before and during photostimulation in the shuttling mice.

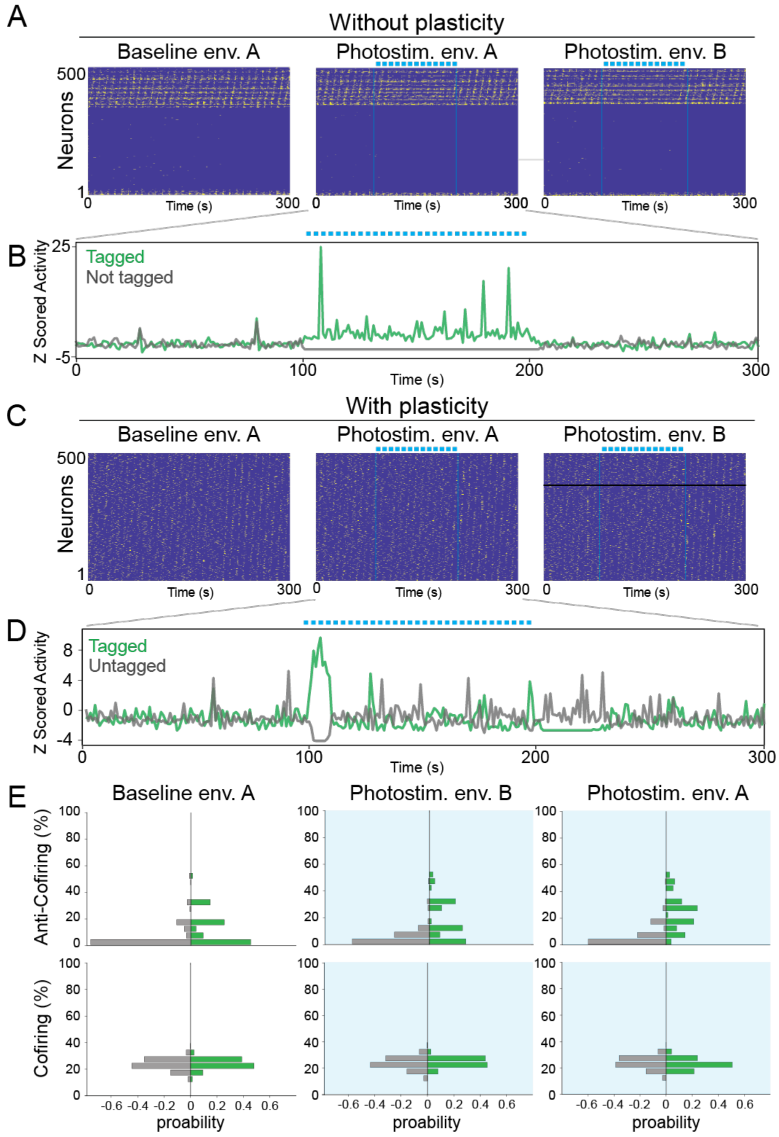


Figure S7. Network simulations suggest that anti-cofiring consequences depend on the concordance between tagging and stimulation conditions and plasticity. A) Rasters of simulated spikes (yellow) without plasticity in two environments. During baseline in environment 1 (left) the top 15% of active units were tagged. The corresponding rasters during photostimulation of the tagged units in environment 1 (middle) and in a distinct environment 2 (right) are shown. B) Normalized population spiking activity during the photostimulation of the tagged (green) and not-tagged (gray) units .C and D) The same as A and B but spike timing-dependent plasticity was enabled during all simulations. Note the initial perturbation of the network in the early stage of stimulation, followed by firing rate homeostasis. E) Anti-cofiring power (proportion of pairs with t < - 0.07) and F) cofiring power of the tagged vs. not-tagged cells in each condition.

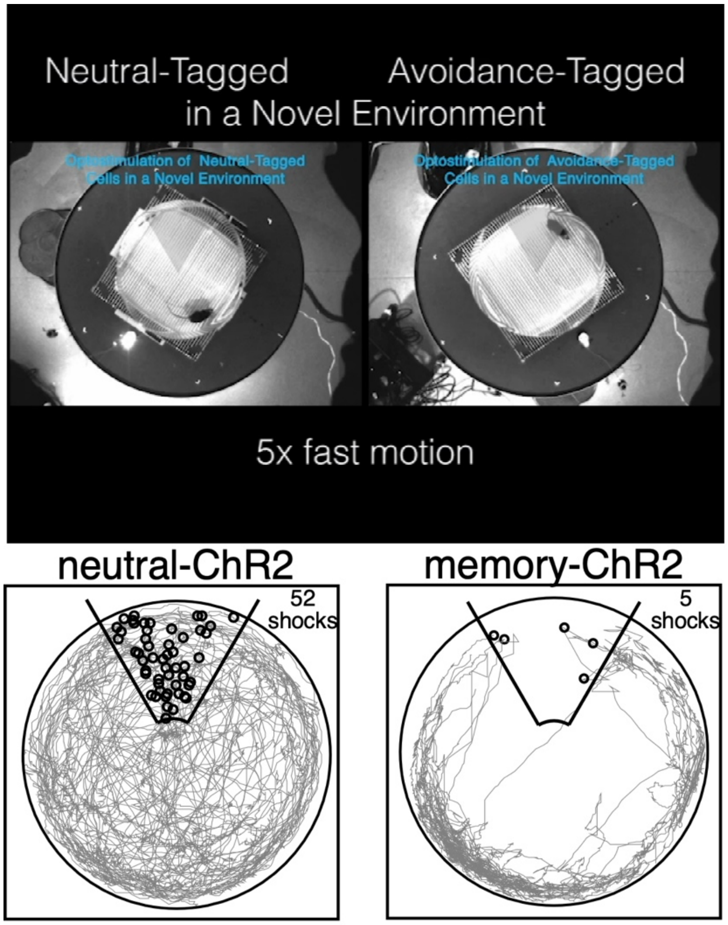


Video S1. Video still (top) from a video during which conditioned active place avoidance behavior is elicited by photostimulation of memory-tagged but not neutral-tagged neurons, while the mice behave in a completely novel environment in which they have never been before and in which they have never experienced shock. (bottom) Tracking from the video illustrating the exploratory behavior of the neutral-tagged mouse (left) and the active place avoidance of the memory-tagged mouse (right). The location of a 60° shock zone is drawn, centered at 12 o'clock along with the positions (small gray circles) of the mice where they would have been shocked if that shock zone had existed.

### METHODS

#### Ethics Approval

All work with mice was approved by the New York University Animal Welfare Committee (UAWC) Protocol ID: 15-1459.

#### Animals

The mice were 3-6 months old. Memory-tagged and neutral-tagged mice were obtained by breeding ArcCreER$^{T2}$(+)[27] × R26R-STOP-floxed-ChR2-eYFP (enhanced yellow fluorescent protein)[45] homozygous female mice with R26R-STOP-floxed-ChR2-eYFP male mice. Control memory-tagged (eYFP) mice were obtained by crossing ArcCreER$^{T2}$(+) × R26R-STOP-floxed-eYFP with R26R-STOP-floxed-eYFP [46,47] homozygous male mice.

**Active place avoidance and control behavior**
Three- to 6-month-old Arc-CreER$^{T2}$-eYFP-ChR2 experimental mice and Arc-CreER$^{T2}$-eYFP control mice (n= 13 total) were trained to form a spatial memory in an Active Place Avoidance task so that memory-activated neurons could be tagged with eYFP-ChR2 or eYFP in the case of controls. The mice were placed in a rotating (1 rpm), 40-cm diameter arena, confined by a 50-cm high transparent cylindrical PETG plastic wall, and their movements were tracked at 30 fps by video tracking from an overhead camera (Tracker, Bio-Signal Group, Acton, MA). The computer defined a shock zone as a 60° annulus-sector in which 20% of the radius was excluded. The shock zone was defined by distal visual landmarks in the room. Upon entering the shock zone, a mouse received a 500-ms, 60-Hz, 0.2-mA foot-shock delivered across parallel stainless-steel rods. A control behavior was used for neutral tagging, where cells were tagged while the mouse explored a familiar neutral environment. Thus all, mice were also exposed to a second, unconditioned rotating arena with a similar cylindrical wall. The parallel-rod floor was covered by clear PETG plastic and there was a distinct scent. The use of one arena for place avoidance and the other for control exploration was counter-balanced across mice. The timeseries of tracked locations was analyzed offline using TrackAnalysis (Bio-Signal Group). Analyses pertaining to the Rayleigh vector of the animal's trajectory were performed and visualized using custom MATLAB R2020b (MathWorks) code.

**Shuttling behavior on a circular track**
Four Arc-CreER$^{T2}$-eYFP-ChR2 mice of both sexes aged 3-6 months old were trained in a foraging task. The track was a 10-cm wide circular corridor created by two concentric PETG clear plastic walls (60 cm and 80 cm inner and outer diameters, respectively). The corridor contained a PETG plastic wall that prevented the mouse from running continuously around circular track, forcing it to shuttle back and forth. In 20-min daily trials, the animals were trained to run toward and stay (500 ms) on one side of the transparent wall (target zone defined by a 8 cm x 10 cm wedge in software, Tracker, Bio-Signal Group, Acton, MA) to activate the automatic delivery of a reward pellet on the opposite side of the wall (20-mg vanilla pellets, Dustless Precision Pellets, Bio-Serv, Flemington, NJ). The animals had to run the length of the circular track and then back to the start to obtain the reward.

**Experimental Design** (Tagging during avoidance behavior)
All animals were implanted bilaterally with 6.4-mm long,1.25-mm OD, 230-µm ID ceramic ferrules (CFLC230-10 Thorlabs, Newton, NJ). The target was above the dorsal hippocampus (-1.9 mm AP, 1.2 mm ML, 1.2 DV). Behavioral training started one week after the surgery.

Isolation housing was used to avoid tagging neurons due to spurious activity unrelated to the memory and control tasks. All mice were housed in a 78 x 52 x 52 cm customized, ventilated, sound-attenuated chamber with a 12/12h light-dark cycle (Lafayette Instruments). The isolation box was within 1 to 3 m of the two rotating arenas. The mice were moved from the vivarium to the isolation box 48 h before, and then returned to the vivarium 48 h after the TAM injection to tag neurons.

After a 30-min habituation trial in each of the memory and neutral environments, each day for four days, the animals were given three 10-min and one 10-min daily training trials in the conditioned and unconditioned environments, respectively. After day two training, the mice were moved to the isolation box. On day five, all animals were given an intraperitoneal injection of 90 mg/kg 4OH-Tamoxifen (TAM) (Sigma) dissolved in corn oil (Sigma) and 100% ethanol either. The mouse was returned to its cage in the isolation box and 45 minutes afterwards, it was placed either on the conditioned or unconditioned rotating arena.

**Experimental Design** (Tagging during shuttling behavior)
After a week of training (one 20-min trial per day for 7 days) the animals were injected with 4OH-TAM 45 min before performing the shuttling task for 30 min, with the intention of tagging active neurons. The animals were housed in a sound-attenuated chamber 1.5 m from the shuttling environment for 48 h before and 48 h after the injection to acclimate them to the environment and reduce spurious tagging when TAM was administered[26]. Three to five days after the injection the animals were implanted with Neuropixel 1.0 probes attached to a 200-µm diameter optic fiber (FT200UMT, Thorlabs Newton, NJ). The target was above the dorsal hippocampus (relative to Bregma -1.9 mm AP, 1.2 mm ML, -1.2 DV). Behavioral training started one week after the surgery.

**Photostimulation**
Two weeks after the TAM injection to tag neurons with ChR2. The effects of photostimulation were assessed either under urethane anesthesia, during head-fixation while running on disk, or during free exploration of the unconditioned arena in which the mouse was never shocked and in which it did not avoid. The optogenetic stimulus was a series of 473 nm wavelength laser pulses (OEM laser Systems, UT). The pulse timing and delivery was triggered by dacqUSB software controlling DI/O hardware (Axona Ltd). The blue light pulses were 15 ms long, 5 mW, and delivered at either 4 Hz or 10 Hz in different mice.

**Single unit electrophysiological recordings under anesthesia**
Single-unit action potential responses to photostimulation were assessed under urethane anesthesia, a condition with hippocampal network activity characteristically reduced ongoing neural activity and excitation-inhibition coupling[41]. Two mice were anaesthetized with 1.20 g $kg^{-1}$ urethane delivered in three intraperitoneal injections spaced 15 min apart, and then placed in a Kopf stereotaxic frame. A craniotomy was drilled and a 16-channel silicon probe electrode (Neuronexus, A1×16-3 mm-50-703) fused to a 200-µm diameter optic fiber attached (Thorlabs, Newton, NJ) was slowly lowered into the dorsal hippocampus (-1.9 mm AP, 1.2 mm ML, 1.4 DV). Using commercial hardware and software (dacqUSB, Axona Ltd., St. Albans, U.K.), electrophysiological signals were amplified 5000 - 7000 times, filtered 300 Hz – 7 kHz and digitized at 48 kHz to record action potentials.

Photostimulation consisted of 15 ms pulses delivered at 10 Hz and 1 mW, 5 mW, or 7 mW. There was no statistically significant differences between the three intensities, so data corresponding to 1 mW stimulation are reported. Action potentials were recorded during a 30-min baseline period and a 30-min photostimulation period. Single-unit action potential waveforms were isolated offline using WClust (https://github.com/FentonLab). Single units were only considered if they were sufficiently well discriminated using objective criteria based on *IsoI* estimates of isolation quality that were above 4 bits [48].

**Single unit electrophysiological recordings in awake head-fixed mice**

For head-fixed recordings 8 mice were anesthetized with 1.5% isoflurane anesthesia and implanted with a titanium head-plate (H.E. Parmer Company, Nashville, TN) using dental cement (Metabond, Parkell, NY) and cyanoacrylate glue (ZAP, Pacer Technology, Ontario, CA). The exposed skull was covered with KwikSil (World Precision Instruments, Sarasota, FL). After a 1-week recovery, a second surgery was performed to drill a craniotomy above the dorsal hippocampus. The craniotomy was covered with Kwik-Sil. The mouse was then placed in a 20-cm diameter, 0.3-cm thick acrylic disk that could rotate on center (Delvie's Plastics, South Salt Lake, UT) and its head was fixed by clamping the titanium head plate to a post. The Kwik-Sil plug was removed and a 200-µm diameter optic fiber (Thorlabs, Newton, NJ) fused to a four-shank, 32-channel silicon probe (Buzsaki32-H32_21mm, NeuroNexus, Ann Arbor, MI) was slowly lowered into the dorsal hippocampus (relative to Bregma -1.9 mm AP, 1.2 mm ML, 1.4 DV), using a Kopf stereotaxic arm. Recordings consisted of a 10-min baseline period, followed by a 10-min photostimulation period (473 nm, 15 ms, 5 mW, 4 Hz or 10 Hz stimulus trains, OEM Laser Systems, UT, externally triggered by dacqUSB, Axona Ltd., St. Albans, UK). A 10-min post-photostimulation recording followed immediately. Data were digitized at 30 kHz, filtered at 300 Hz - 6 kHz, and collected using an OpenEphys (https://open-ephys.org) acquisition system and a 32-ch Intan headstage (Intan Technologies, UT). Single units were automatically sorted using the open-source algorithm Kilosort2 [49], then manually sorted with the open-source Python library Phy (https://github.com/cortex-lab/phy). All subsequent single-unit data analyses were also performed using custom MATLAB code (MathWorks, Natick, MA). Cells with mean firing rate of >5 AP/s during the 10-min baseline and with narrow (< 420 µs) waveforms were considered putative interneurons.

**Single unit electrophysiological recordings in freely-behaving mice**

All animals were anesthetized with 1.5% isoflurane and implanted with a Neuropixel 1.0 probe (Imec) attached to a 200-µm diameter optic fiber (Thorlabs, Newton, NJ) in dorsal hippocampus (relative to Bregma -1.9 mm AP, 1.2 mm ML, 1.2 DV). The probe was attached to a custom 3-D-printed apparatus and mounted on the head with bone screws and dental cement. One week after surgery single-unit activity was collected using commercial hardware (PXIe Control System, Imec, with National Instruments chassis) and software (SpikeGLX, Bill Karsh, https://github.com/billkarsh/SpikeGLX) while the animals performed the foraging task. Single units were isolated using Kilosort 2.5 and Phy for manual curation as described above.

**Photostimulation during free exploration of the unconditioned arena**

Thirteen mice were tested to determine whether photostimulation of the tagged cells would elicit conditioned avoidance in the environment where the mouse never experienced shock. Mice from the memory-ChR2, memory-eYFP, and neutral-ChR2 groups were placed in the unconditioned environment and photostimulated for 10 min at either 4 Hz or 10 Hz.

**Statistical analyses**

All statistical analyses were performed using JMP Pro 16.0.0 (SAS, Cary, NC). Statistical significance was assessed by t tests or multiple factor comparisons performed by ANOVA, with or without repeated measures, followed by Tukey post hoc tests as appropriate. Statistical comparisons between proportions of binomial outcomes were performed using the test of proportions. The alpha threshold for significance was set at 0.05.

**Neural network simulations**
The Excitatory-Inhibitory (E-I) Recurrent Network used in Fig.1 and described in detail [22,50] was adapted to model optogenetic stimulation. Briefly, the recurrent neural network model of 500 excitatory and 50 inhibitory units with all-to-all connections receives random inputs from a circular track that the agent moves along. The E-I, I-E, and I-I connection weights adjusted according to spike timing dependent plasticity (STDP) rules without supervision, which generates activity representations of the agent's current location on the track. In addition to the positional inputs, the recurrent excitatory inputs, and the recurrent inhibitory inputs, a random 20% subset of "tagged" E cells received an additional input meant to simulate optogenetic stimulation. The optogenetic stimulation was excitation of 50% of the spiking threshold for 14 ms, every 100 ms during the STIM ON conditions, which modeled the patterned photostimulation onto randomly tagged units.

*Distinctiveness*: Normalized firing rates were calculated to generate spatial firing rate maps by dividing the number of spikes at a given location by the time spent in that location. The firing rate map across all E cells and positions were concatenated and normalized by the maximum firing rate at each position across cells. The distinctiveness measure, ranging from 0 to 1, was then calculated as the average rate across all positions and E cells in the recurrent network.

Three simulations were run to model optogenetic tagging and stimulation, with randomly initialized inputs defining one of two environments. Environment 1 was a baseline condition to find the highest firing cells. The top 15% cells with highest mean rate were designated the tagged cells). Then, using the same Environment 1 inputs or a different randomly initialized inputs (Environment 2), the tagged cells were driven by an additional "external" input (causing EPSPs that are 25% of the potential difference between the reset potential and the threshold potential). The additional input consisted of pulses of stimulation (14 ms pulses that onset every 100ms) to model the experimental photostimulation.

**Immunohistochemistry**
In order to confirm cell tagging with Channelrhodopsin as done previously[26], brains were fixed by cardiac perfusion and placed in ice-cold 4% paraformaldehyde in 0.1 M phosphate buffer overnight (PB, pH 7.4) and for 24 h in each of 10%, 20%, and 30% sucrose solution for cryoprotection. The brains were then cut in 40-µm slices with a Leica cryostat. The slices were then washed in Tris-buffered saline (TBS) three times for 5 min. After blocking the slices in 0.05 M TBS containing 5% normal goat serum (NGS) and 0.25% Triton X-100 for 1 h at room temperature, the sections were incubated in the primary antibody, chicken anti-GFP (Abcam, ab13970) diluted to 1:500 in the same TBS/NGS/Triton blocking solution and incubated at 4 °C for 48 h. After washing the slices three times for 20 min in TBS, they were incubated for 2 h in the secondary antibody (Alexa Fluor 488-conjugated goat anti-chicken IgY (H+L), Thermo Fisher, A32931), diluted to 1:200 in the TBS/NGS/Triton blocking solution, washed again three times for 20 min in TBS and mounted on Superfrost Plus slides (Fisherbrand) and covered using ProLong Gold Antifade (Thermo Fisher, P36930) mounting medium with DAPI. Imaging was

performed using a Leica TCS SP5 confocal microscope, with a 40x oil objective lens. After acquisition, images were deconvolved using Huygens Professional software using classic maximum likelihood estimations (Scientific Volume Imaging, Hilversum, Netherlands) and processed with ImageJ/Fiji software (US National Institutes of Health). Quantification of eYFP-positive cells was performed manually.
